# Fusicatenibacter Is Associated with Kefir Drinking

**DOI:** 10.1101/218313

**Authors:** Richard Sprague

## Abstract

Daily 16s rRNA-based microbiome sampling reveals that consumption of the fermented drink, kefir, is associated with a previously-unexplored genus *Fusicatenibacter* of the *Firmicutes* phylum within family *Lachnospiraceae.*

## Introduction

Kefir is a fermented milk drink produced by the action of bacteria and yeasts and believed to have medicinal uses. A rigorous microbial analysis by Walsh et al. (2016)^1^ recently showed precisely which microbes are present in kefir, at various stages in the fermentation process. (See Figure ŒŒ). The grains themselves contain a combination of lactic acid bacteria *(Lactobacillus, Lactococcus, Leuconostoc),* acetic acid bacteria *(Acetobacter),* and yeast, clumped together with casein (milk proteins) and complex sugars in a matrix of a unique polysaccharide called Kefiran. The nutritional content apparently varies depending on fermentation time and other factors.^2^

Numerous studies indicate that regular kefir drinking has positive effects on health, and it is reasonable to assume that its known bacteriological properties would affect the gut microbiome, but we are unaware of previous research that conclusively demonstrates that microbes in kefir make it successfully through the acidic environment of the stomach.

Although similar doubts have been expressed about another fermented dairy product, yogurt, careful research has shown several microbial strains that pass through the body successfully. (Uyeno, Sekiguchi, and Kamagata (2008)) Furthermore, several of the microbes apparently persist in the gut and can be observed a full 28 days after consumption.

We were interested to know if the same is true of kefir, and how it might alter the gut microbiome on a daily basis. We sequenced the 16S rRNA gene in 500 near-daily samples of the microbiome of a single subject, a 50-year-old male in excellent health. Replicating the experiment in David et al. (2014), we carefully tracked diet, sleep, location, activity, and other variables. Most samples were from gut, but bimonthly samples were regularly taken of skin, nose, and mouth as well.

Because we had several hundred days worth of daily microbiome sampling before the subject first encountered kefir, we also wanted to find if any new microbes appeared (or disappeared) as a result. Finally, by continuing to test long after the kefir consumption began, we were able to see how long any such microbes remain in the gut.

## Results

We found that kefir consumption was associated with a clear change in the abundance of several organisms, including one, *Lactococcus,* whose presence could be confirmed in the drink as well. To our surprise, we also found at least one new organism, a novel one that had not been observed in hundreds of previous samples taken from the same subject. Furthermore, the new organism, *Fusicatenibacter,* appears to remain in the gut after ending the keir consumption, indicating a persistent alteration of the gut microbiome.

The high levels of *Lactococcus* in the subject’s gut was not unexpected. Previous studies have shown it to inhabit kefir drinks^3^ and we also found it as the main genus of microbe found in our kefir samples as well. (see Table 1) We found that, indeed, it survives passage through the stomach and presents in the subject’s gut in high quantities. Because we have the subject’s near-daily gut microbiome records for more than one year prior to his first consuming kefir (See Figure 1), we confidentally attribute the newly-found abundance to the drink.

**Table 1:** Sequenced abundances found in the kefir drink before consumption.

|  | Kefir (%) |
| --- | --- |
| Lactococcus | 96.06 |
| Leuconostoc | 3.02 |
| Lactobacillus | 0.22 |
| Faecalibacterium | 0.14 |
| Roseburia | 0.06 |

**Figure 1:**
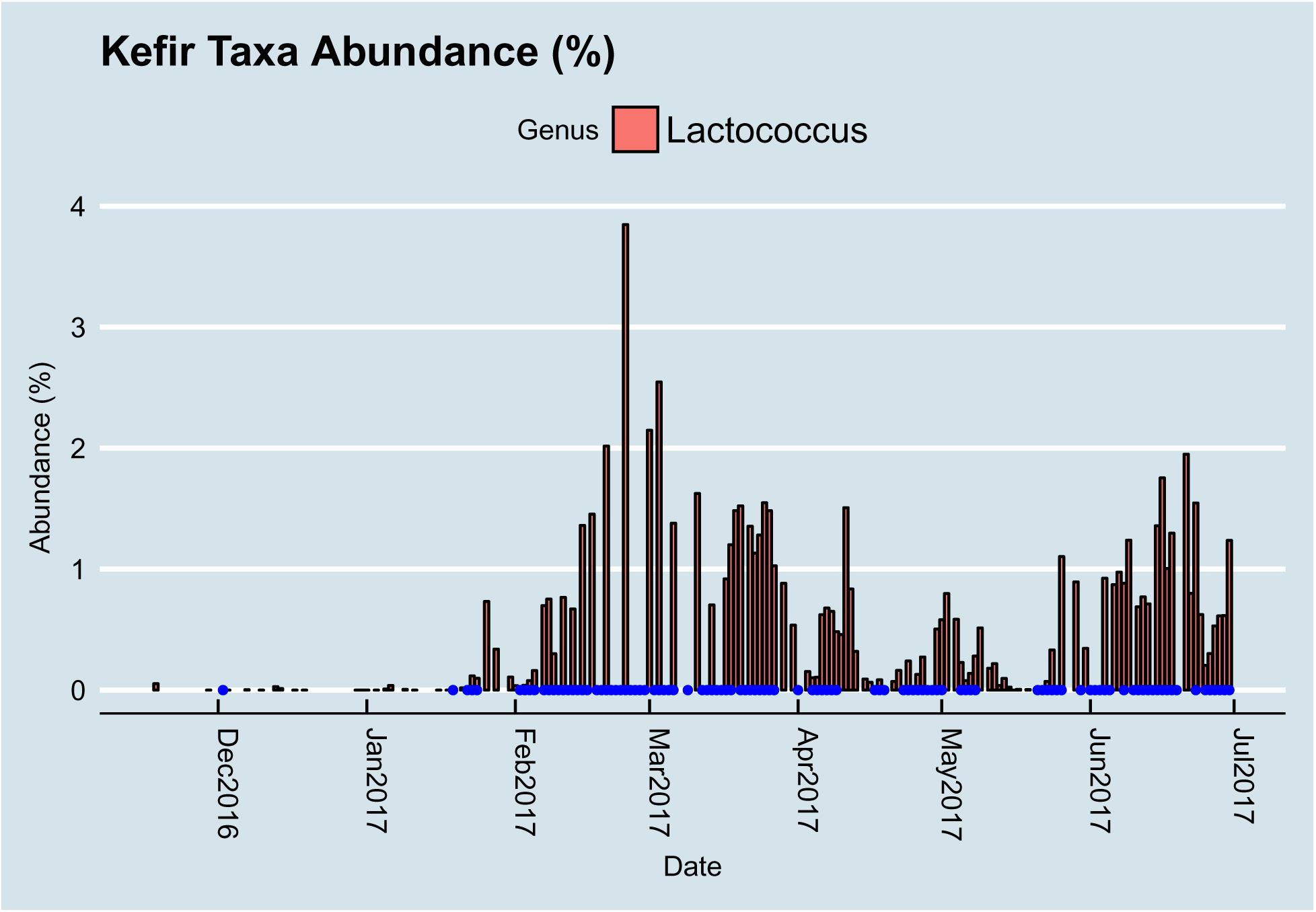
Abundance levels of Genus *Lactococcus* in a 55-year-old male subject’s gut. Kefir was consumed only on the dates indicated in blue. Samples were taken near-daily throughout the period, so abundance levels are zero unless otherwise indicated. We note that levels seem to dip when on days when the kefir is not consumed, such as during trips out of town in mid-April and mid-May.

We also spotted a new new microbe, *Fusicatenibactor* that appears to exactly trace the kefir consumption. (Figure 2). A gram-stain-positive, obligately anaerobic, non-motile, non-spore-forming, spindle-shaped bacterium, our literature search revealed nothing more of interest since its isolation in 2013^4^. Although it is seen regularly in the human gut, we are unaware of any reports of its connection to diet. We believe this is the first reported instance of its association with a specific type of food.

**Figure 2:**
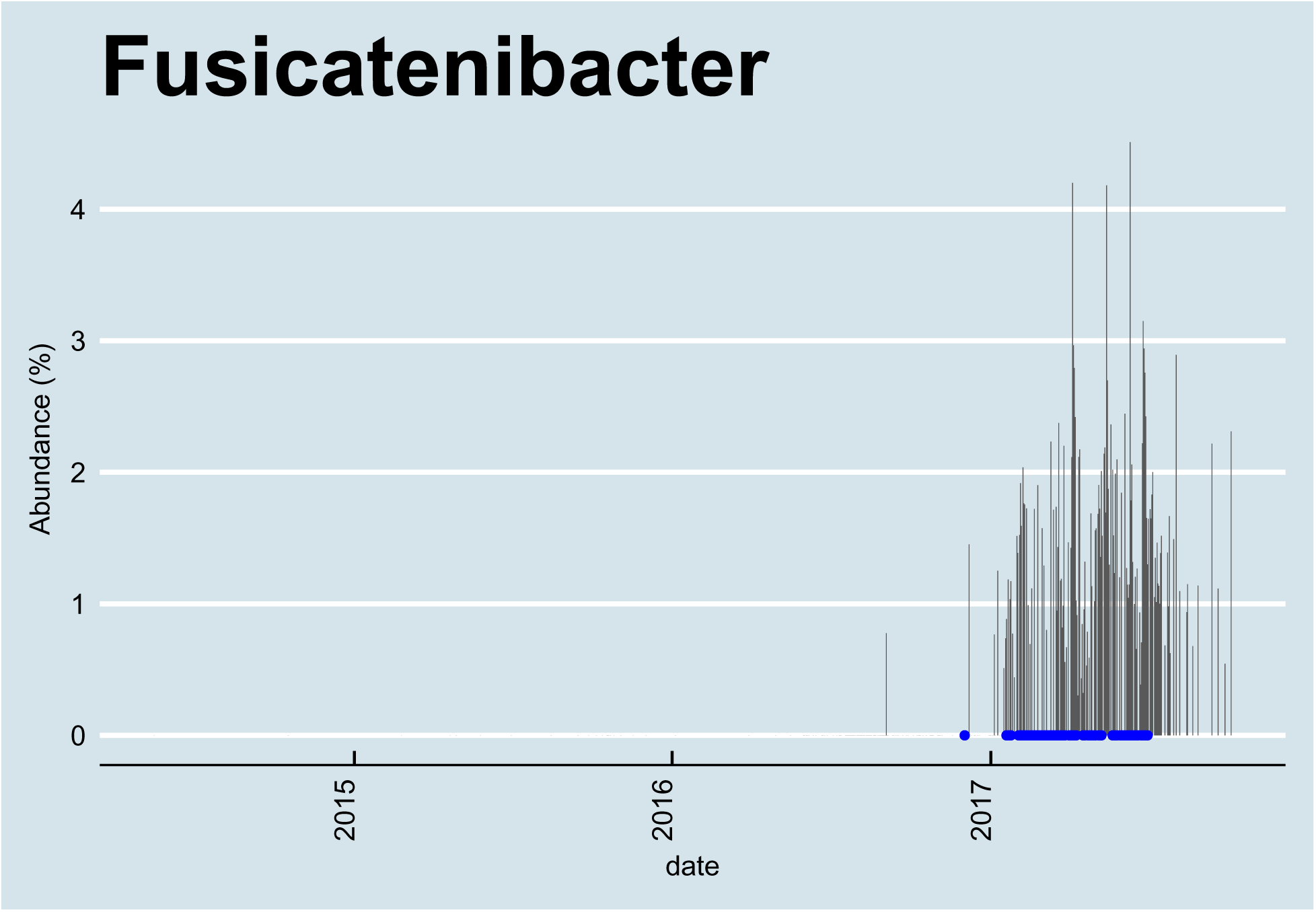
*Fusicatenibacter* is found at high abundance after drinking kefir. This chart shows abundance levels were zero since testing began more than two years previously.

## Discussion

Although we were pleased to see that one of the microbes in the original drink made it through the body and could be found in the gut, this result alone merely confirms what intuition would suggest: microbes that go in the mouth can successfully navigate the entire gastrointesintal tract. Other researchers have reported similar persistence of the same microbe *Lactococcus* after yogurt consumption, another fermented drink. Interestingly, our subject, an occasional yogurt eater, showed virtually none of this microbe in the years of measurement before drinking kefir. We speculate that there may be something uniquely robust about the particular species of *Lactococcus* found in this sample, one that may not be found in the commercially-available yogurt previously consumed by the subject.

It is interesting to note that the subject was drinking homemade keir, fermented overnight in his kitchen, and thus exposed to the same environmental microbes that would have surrounded the subject himself. We hypothesize that the known high variance in microbial environments may play a significant role in which microbes appear in the gut. Commercially-purchased kefir is produced in sanitized industrial environments which, while enabling a consistent product and protective against pathogens, may inevitably result in differences in microbial strains.

We do not understand why a novel microbe, *Fusicatenibacter* would appear in the gut in such large quantities immediately after the first drink. We confirmed with the lab that this microbe was unlikely to result from contamination. Although it had not been found in this subject previously, the lab reports that it is found regularly in samples from other people. Analysis of the plates on which the subject’s samples were processed indicated no irregularities; in fact, the wells directly adjacent to the subject’s sample did not show any of this microbe, though that was present in other samples processed in the same run.

A literature search reveals nothing of clinical or other apparent interest about this microbe, a Clostridium that appears within the family *Lachnospiraceae* of phylum *Firmicutes.* We can find no apparent link to health or other conditions documented by other projects. Since it persists and makes up from 1-4% of the subject’s post-kefir microbiome, we think it must have found a role in the microbial ecosystem.

Note that the subject remained in excellent health before and after the kefir consumption. We could detect no significant differences in blood chemistry or other quantitative health metrics. A review of his activity, sleep, and diet reveals no other significant differences that might compound the microbiome changes that occurred after beginning keir.

## Methods

Samples were collected on a daily basis, following instructions from commercially-available kits from uBiome, Inc. Fecal samples were lightly mixed and swabbed throughout to lessen distribution anamolies within the sample. The swabs were stirred into a lysis buffer and then transported at room temperatures to the uBiome lab. Published accounts of the uBiome protocol indiciate that genomic DNA was extracted by a liquid-handling robot, amplified up to 30 times using PCR, with primers inserted at the V4 subunit of the rRNA gene ((515F: GTGCCAGCMGCCGCGGTAA and 806R: GGACTACHVGGGTWTCTAAT) using Illumina NextSeq platform rendering 2 × 150bp pair-end sequences.

Samples were barcoded with a unique combination of forward and reverse indexes allowing for simultaneous processing of multiple samples. De-multiplexing of samples was performed using Illumina’s BCL2FASTQ algorithm. Acquired reads were filtered using an average Q-score > 30. Primers and any leading bases were subsequently trimmed from the reads, and forward and reverse reads were appended together. To effectively cluster real biological sequences and to identify reads that contain errors as a product of sequencing, reads were clustered using the Swarm algorithm (Mahé et al. 2014) using a distance of 1 nucleotide. The most abundant sequence per cluster was considered the real biological sequence and was assigned the count of all reads in the cluster.

The representative reads from all clusters were subjected to chimera removal using the VCHIME algorithm (Rognes et al. 2016). Reads passing all above filters were aligned using an in-house uBiome database of 16S sequences derived from the NCBI-nr database (Benson et al. 2013). Decreasing sequence identities were used to map reads to different taxonomic rankings: > 97% sequence identity was used for the assignment to a species, > 95% sequence identity for the assignment to a genus, > 90% for assignment to a family, > 85% for assignment to an order, > 80% for assignment to a class, and > 77% for assignment to a phylum. The relative abundance of each taxonomic group was calculated by dividing the abundance of the taxonomic group to all sequences that map to any sequence in the bacterial domain.

Bionformatics was performed in R using Phyloseq (McMurdie and Holmes (2013)). Source code is available on our Github page.

## Conclusions

Interesting associations with microbes of unknown function are commonly found in all microbiome experiments, and the volume of data collected ensures that some "statistically-significant" results will be found simply due to random chance. Although the function of *Fusicatenibacter* is unknown, the association in this subject was so pronounced, and so clearly associated with the start of kefir drinking that we felt other researchers may benefit from learning about our experience. Despite the n=1 oddity of this experiment, we have seen the organism in other kefir-drinking subjects, and we present this work in the hopes that it may be useful others working to understand the effect of kefir and other fermented drinks on human health.

## Footnotes

1 also see a 2-minute Youtube presentation

2 Otles and Cagindi (2003) and http://files.cienciapatodos.webnode.pt/200000022-79ffe7af9e/Kefir.pdf

3 Leite et al. (2013)

4 Takada et al. (2013)

